# Predicting Obstetric and Non-obstetric Diagnoses Co-occurrences during Pregnancy

**DOI:** 10.64898/2026.02.06.704385

**Authors:** Akash Singh, Samuel Infante, Seungbae Kim, Anowarul Kabir

## Abstract

Pregnancy care often involves simultaneous obstetric and other medical conditions, but their co-occurrence patterns are rarely modeled explicitly in a systematic, network-based approach. In this work, we formulate obstetric and non-obstetric diagnoses co-occurrences as a link prediction problem on a diagnosis-level homogeneous graph constructed from pregnancy encounters. Diagnoses are represented as nodes connected by co-occurrence edges, with node features capturing graph structure and demographic statistics^3^.

We address this challenge by leveraging collected electronic health records data and study several standalone and hybrid graph neural network (GNN) architectures, including GCN, GAT, GraphSAGE, and three hybrid encoders that combine complementary aggregation mechanisms, namely GCN+GraphSAGE, GCN+GAT, and GAT+GraphSAGE. All models used consistent train-validation-test splits and are evaluated on 5- fold cross-validation sets. Among standalone models, GraphSAGE achieved the strongest performance, whereas hybrid GraphSAGE-based models (GCN+GraphSAGE and GAT+GraphSAGE) are best performers. The GCN+GraphSAGE hybrid, reaching an AUROC and AUPRC of approximately 0.90, consistently outperformed all other architectures. Further analysis of top-ranked predicted links revealed clinically plausible associations between pregnancy stage and risk-related diagnoses and common endocrine, metabolic, and hematological conditions. These findings indicate that graph-based link prediction may effectively prioritize obstetric and non-obstetric diagnosis pairs, providing a scalable framework for identifying clinically meaningful comorbidity patterns. They may further support hypothesis generation and downstream obstetric risk stratification efforts.

**Availability:** All codes including data preparation scripts, training and validation recipes, and experimental configurations are available at: https://github.com/kabir-ai2bio-lab/ob-nonob-diagnoses-cooccurrences.

## 1 Introduction

Pregnancy is a high-stakes period where preexisting chronic conditions, acute complications, and social risk factors interact in a complex way. Maternal morbidity and mortality are often diagnosed by conditions related to pregnancy or postpartum period, known as obstetric diagnoses. However, combinations of preexisting or developing medical or surgical non-obstetric conditions, such as cardiovascular, metabolic, hypertension and/or anxiety disorders, may also contribute to the ill cause simultaneously. Since clinicians often focus on obstetric diagnoses, automatic diagnosis of non-obstetric and obstetric co-occurrences during pregnancy is of great interest for risk assessment, pregnancy care and fetal outcomes.

There is a growing body of interest in developing machine and deep learning based methods for risk prediction [18] and clinical diagnosis [6,25]. This is due to the availability of high dimensional patients’ electronic health records (EHRs) that document patients’ symptoms, prescriptions, clinical notes, and medical images which is heterogeneous and longitudinal in nature [14,22,20]. Despite this high-volume and rich information present in many EHRs, almost no research has been conducted concentrating both obstetric and non-obstetric conditions co-occurrences focusing on pregnant morbidity and mortality. Recent systematic review of ML models focuses on obstetric outcomes (e.g., preeclampsia) [12,17,7]. Meredith et al. [11] highlight that including preexisting co-occurring medical conditions (multimorbidity) improves predictive performance for severe preeclampsia. Several works have concentrated on maternal and adverse birth outcomes for obstetric diagnoses, broadly summarized in [13,1,2]. Therefore, to address this gap in pregnancy care, we propose graph neural network (GNN) based frameworks and evaluate multiple architectures to predict obstetric and non-obstetric diagnosis co-occurrences during pregnancy, essentially providing a benchmark for future development.

We formulate the problem as a network of co-occurring diagnoses rather than isolated events, since EHRs store these obstetric and non-obstetric conditions as coded diagnoses across temporal encounters of the same patient. We ask *which obstetric and non-obstetric diagnosis pairs are more likely to co-occur in future pregnancy encounters*? We construct a diagnosis-level graph from pregnancy-related encounters. Each node represents a diagnosis concept, edges represent co-occurrence of two diagnoses within the same pregnancy visit. Then, the nodes are enriched with structural features from the graph plus demographic statistics, such as average patient age at which the diagnosis appears in pregnancy encounters. We then formulate a link prediction problem, given the current pregnancy diagnosis graph and node features. To solve this problem, we systematically applied and studied various existing GNNs and hybrid architectures for robust and explainable predictive modeling towards co-occurrence diagnoses.

Several studies have applied GNNs to EHR-like data. Zhu et al. [26] proposed a variationally regularized graph-based representation learning framework that uses GCN layers over a medical concept graph to learn patient embeddings for downstream prediction tasks. Cho et al. [3] built a heterogeneous graph of patients and medical entities and used HinSAGE, a variant of GraphSAGE for heterogeneous graphs, to learn patient representations that improved prognostic prediction compared with non-graph baselines. More recent heterogeneous GNN frameworks, such as GraphEHR, also use neighborhood-sampling-style message passing to predict multiple diseases from hospital EHR data [9]. Graph attention networks (GATs) have been explored for several clinical tasks. GRAM introduced an attention mechanism over a medical ontology tree to compute concept representations for disease prediction, highlighting that attention over graph neighbors can improve interpretability [4]. HealthGAT proposed a hierarchical GAT architecture that learns embeddings for medical codes and visits from EHR graphs and showed gains on node classification and readmission prediction tasks [15]. GAT variants have also been used together with ontology trees for medication recommendation, again showing that attention over clinically meaningful graph structure can enhance performance [24].

There is growing interest in graph transformers for EHR. Hypergraph Transformers for EHR-based clinical predictions use hyperedge-based self-attention over sets of medical codes within visits, and achieve strong results on readmission and mortality prediction tasks [23]. Poulain et al. [16] proposed a graphbased time-aware transformer that builds visit embeddings with a graph module over medical codes and then applies a BERT-style model to patient timelines, improving prediction of future diagnoses. These works suggest that transformer-style attention over graph-structured EHR data can capture complex dependencies beyond what local message-passing GNNs see. The Graph Convolutional Transformer (GCT) combines GCN layers over a diagnosis code graph with a transformer over visit sequences, and shows that explicitly learning the graphical structure of EHR codes improves the prediction of future clinical events [5].

Although the application of GNNs to EHR data is not new, recent works [18,14] mentioned that translating EHRs into meaningful insights is challenging due to the heterogeneity, high-dimensionality, poor-quality, sparse and multimodality present in the data. Unlike temporal or heterogeneous EHR graph models that learn representations at the patient or visit level across longitudinal data, our approach deliberately adopts a diagnosis-centric homogeneous graph abstraction. Pregnancy encounters are collapsed into a co-occurrence network to capture structural comorbidity patterns that are often diluted in patient sequence models. This design complements time-aware EHR graph approaches by offering an interpretable view of interactions between obstetric and non-obstetric diagnoses. In this work, we create and propose a novel benchmark of standalone and hybrid GNN architectures that focuses on predicting the likelihood of obstetric and non-obstetric diagnoses co-occurrence in pregnancy encounters, rather than patient-level outcome prediction or general diagnosis forecasting. We leverage a diagnosis co-occurrence graph, systematically evaluate multiple GNN families, including GCN [10], GraphSAGE [8], GAT [19], and hybrid approaches, in order to identify which architecture choice affects the ability to predict clinically meaningful co-morbidity links in pregnancy care. Our proposed model can enable non-obstetric conditions to appear together with pregnancy-related diagnoses and may act as comorbidity hubs. With our reliable and rigorously evaluated approach, we aim to support earlier identification of clinically relevant comorbidity patterns that may inform downstream risk stratification, targeted monitoring, and counseling for patients at risk of multi-morbid pregnancy courses. However, this work is designed to model diagnosis-level co-occurrence patterns within pregnancy-related encounters, rather than to perform patient-level outcome prediction or real-time clinical decision support. Accordingly, the proposed framework is most applicable to retrospective EHR analysis for hypothesis generation, cohort characterization, and risk pattern discovery. It is not intended to replace clinician judgment or establish causal relationships.

## 2 Problem Setup

To identify the co-occurring obstetric and non-obstetric diagnoses, we define a homogeneous undirected diagnosis graph, *G* = (*V, E*) where *V* defines the set of diagnoses concepts as graph nodes that often appear during pregnancy encounters in addition with preexisting conditions, and *E* denotes the set of edges or relations between various diagnoses concepts. Next, we define the edges’ weight and compute feature vector for each node, described as follows.

### 2.1 Defining Relation among Diagnosis Concepts as Edges

Edges encode the co-occurrence of diagnoses within pregnancy encounters. For each pregnancy visit *r*, we collect the set of unique diagnoses

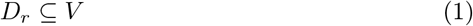

For each unordered pair of diagnosis, (*i, j*) ⊆ *D*_*r*_, we compute a visit level co-occurrence counter by incrementing by one each time.

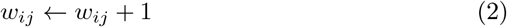

After processing all pregnancy visits, we define an undirected edge (*i, j*) ∈ *E* if and only if *w*_*ij*_ ≥ 2. Each edge therefore has an integer weight (*w*_*ij*_) equal to the number of pregnancy encounters in which diagnoses *i* and *j* co-occur. Additionally, we define an auxiliary edge distance for path-based measures and community detection

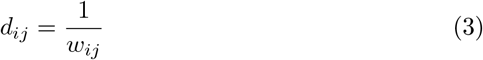

such that paths through strongly co-occurring pairs are shorter.

### 2.2 Defining Diagnosis Concepts as Nodes

Each node *v* ∈ *V* corresponds to a unique diagnosis concept that appears in the dataset by at least ten pregnancy encounters. We define the node features **x**_*v*_ ∈ ℝ^9^ by considering both graph structural and patient demographic information, discussed as follows:

**– Pregnancy degree.** The unweighted degree of *v* in the pregnancy graph,

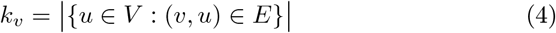

**– Pregnancy co-occurrence strength.** Let *w*_*vu*_ be the edge weight between *v* and *u*. The co-occurrence strength is defined by

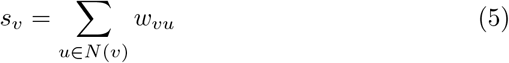

where *N* (*v*) is the neighboring nodes of *v*. This counts how often *v* co-occurs with any other retained diagnosis in pregnancy encounters.

**– Betweenness centrality in the pregnancy graph.** Using the edge distance definition from Eq 3, we compute standard betweenness centrality for each concept

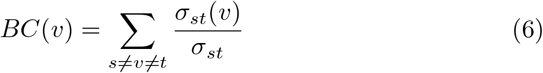

where *σ*_*st*_ is the number of shortest paths between *s* and *t* and *σ*_*st*_(*v*) counts those that pass through *v*. This measures how often *v* lies on shortest co-occurrence paths.

**– Community participation coefficient.** We run Louvain method [13] to extract non-overlapping communities on our defined concept network *G*. Let 𝒞 denotes the set of communities and 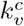 the number of neighbors of *v* that lie in community *c* ∈ 𝒞 . The participation coefficient is computed as

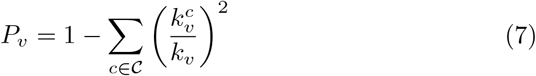

which is high when *v* distributes its edges across many communities and low when it is embedded inside a single community.

**– Cross community neighbor share.** Let *ℓ*(*v*) ∈ {OB, nonOB} be the obstetric label of node *v*. The cross-community neighbor share is

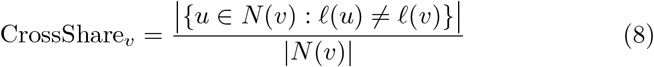

which captures the proportion of neighbors that belong to the opposite class (obstetric vs non-obstetric).

**– Mean age at diagnosis in pregnancy.** The mean age of patients at which diagnosis *v* appears in pregnancy encounters.

**– Age variability at diagnosis in pregnancy.** Standard deviation of patient age when the diagnosis is recorded.

**– Obstetric concept indicator.** Binary feature indicator of whether the diagnosis belongs to the obstetric vocabulary.

**– Bridge neighborhood indicator.** Binary feature equal to 1 if the diagnosis has at least one obstetric neighbor and at least one non-obstetric neighbor.

All continuous features (items 1–7) are standardized globally to have a mean of zero and unit variance. The last two remain binary indicators. This feature design allows the models to combine global graph structure, community connectivity, and demographic information when predicting which potential edges are likely to exist.

### 2.3 Obstetric and Non-obstetric Diagnoses Co-occurrences

We define the prediction task of obstetric (OB) and non-obstetric (nonOB) diagnoses co-occurrences as the link prediction problem over the set of possible obstetric–non-obstetric pairs. Positive edges are the set of observed cross-community edges, defined as

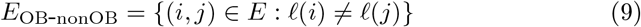

And, the negative edges are between concept pair (*i, j*) such that

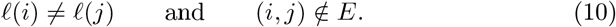

The full graph *G*, including obstetric–obstetric and non-obstetric–non-obstetric edges, provides the structural context used by the GNN encoders.

## 3 Dataset and Preprocessing Steps

We utilize the EHRShot sampled EHR dataset [21] (2, 000 patients in OMOP format) and focused on conditions recorded in the `sampled_condition_occurrence` file table. We kept only rows with non-null `person_id`, `visit_occurrence_id`, and `condition_concept_id`, then created string identifiers for patient, visit, and diagnosis. Empty diagnosis codes were removed. Then we apply several global and local filters.

As a first global filter, we required that a diagnosis concept appear at least 20 times in the full dataset. This removed extremely rare codes that would produce unstable statistics. We also capped the vocabulary to the 2, 000 most frequent diagnoses by global count, although the pregnancy graph became much smaller after subsequent restrictions.

Pregnancy-related concepts were identified in `concept.csv` by searching condition names for a curated list of substrings such as “pregnan”, “obstetric”, “prenatal”, “antepartum”, “postpartum”, “gestational”, “preeclampsia”, and “labor”. The corresponding concept IDs define the set of obstetric diagnoses. A visit was considered a pregnancy encounter if at least one of its diagnoses belonged to this obstetric set. All subsequent graph construction is restricted to these pregnancy encounters.

Within the pregnancy cohort, we applied a second level of filtering. A diagnosis had to appear in at least 10 pregnancy encounters to be retained as a node, and a pair of diagnoses had to co-occur in at least two distinct pregnancy encounters to be kept as an edge. These explicit thresholds balance two goals: removing noisy one-off co-occurrences and keeping enough connectivity for meaningful community structure and centrality measures.

Demographic information was added by joining the filtered condition records with `sampled_person.csv` on `person_id`. We used `year_of_birth` and the condition date (when available) to compute age at each diagnosis event. Implausible ages (negative or greater than 120) were set to missing. For each diagnosis, we then computed the mean and standard deviation of age across all pregnancy encounters where it appeared. Missing age statistics were imputed with the global mean age (for the mean) and zero (for the standard deviation).

All continuous node features were standardized with a global z-score transform before being saved. Every model in the project, including GCN, Graph-SAGE, GAT, and the hybrid architectures, loads exactly this preprocessed graph and uses the same train, validation, and test splits for link prediction.

After all filtering and preprocessing, the pregnancy diagnosis co-occurrence graph used for training contains 279 unique diagnosis concepts as nodes, 5, 285 undirected co-occurrence edges and 9 features per node. Based on the obstetric label of each diagnosis, 124 nodes (44.4 percent) are obstetric, and 155 nodes (55.6 percent) are non-obstetric. The edge set partitions as: (i) Obstetric–non-obstetric edges: 2, 717 edges (51.4 percent), (ii) Obstetric–obstetric edges: 1, 628 edges (30.8 percent), and (iii) Non-obstetric–non-obstetric edges: 940 edges (17.8 percent).

### Data splits

We focus on edges between obstetric and non-obstetric diagnoses:

- Collect all observed cross-community edges and randomly split them into 70% train, 15% validation and 15% test positives.
- For each split, sample an equal number of negative pairs between obstetric and non-obstetric nodes that are not directly connected in the graph.
- Validation and test positive edges are removed from the adjacency matrix used for message passing to avoid label leakage.

The same splits are reused across all architectures.

## 4 Methodology

We perform binary link prediction on obstetric and non-obstetric diagnosis pairs, using observed cross-community edges as positives and randomly sampled un-connected pairs as negatives, with fixed splits reused across all architectures. We consider three GNNs and three hybrid architectures in model development process that follow the same encoder-decoder structure. First, we briefly discuss the GNN architectures that computes the nodes representation from the graph, followed by the decoder as the prediction module.

### 4.1 Graph Convolutional Network (GCN)

The GCN uses three graph convolutional layers with hidden dimension 128, batch normalization, and dropout. At layer *ℓ*, node representations **H**^(*ℓ*)^ are updated by

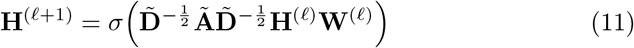

where Ã is the adjacency matrix with self-loops, 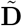 is its degree matrix, **W**^(*ℓ*)^ is a learnable weight matrix and *σ* is a ReLU nonlinearity. This corresponds to a degree-normalized neighborhood average that smooths features over local pregnancy co-occurrence structure.

### 4.2 Graph Attention Network (GAT)

The GAT model uses two attention layers with four heads, exponential linear unit (ELU) activations, batch normalization, and dropout. It replaces uniform aggregation with learned attention weights. For one attention head, coefficients between node *v*, neighbor *u* and the update are

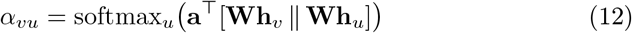

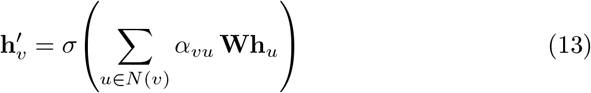

Multiple heads are combined (first by concatenation, then by averaging in the last layer) before the decoder. This allows the model to focus on specific comorbid diagnoses that are more informative for predicting cross-obstetric vs non-obstetric links.

### 4.3 GraphSAGE

The GraphSAGE model also uses three layers with a hidden size of 128, but replaces spectral smoothing with an explicit aggregation of neighbors features. For a node *v* at layer *ℓ*,

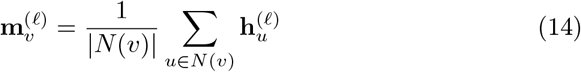

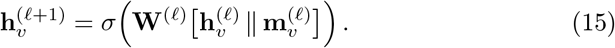

where *N* (*v*) is the set of neighbors and ∥ denotes concatenation. This preserves the node’s own representation while injecting a learned summary of its neighborhood, which can capture heterogeneous local patterns between different diagnosis communities.

### 4.4 Hybrid GCN + GAT

The first hybrid model runs a GCN branch and a GAT branch in parallel on the same graph. The GCN branch applies a stack of GCNConv layers with ReLU and dropout between layers. In contrast, the GAT branch applies a stack of GATConv layers with multi-head attention using a nonlinear transformation through ELU and dropout between layers. Importantly, both branches are configured to produce the same final embedding dimension (`out_channels`), after which layer normalization is applied separately to the GCN and GAT outputs to stabilize their scales before fusion. For each node, the two normalized branch embeddings are then fused either by (i) concatenation followed by a linear projection back to `out_channels` (`projected_concatenation`), or (ii) a learnable gated weighted sum (`gated_sum`) that adaptively balances the contribution of the GCN and GAT representations per node. Conceptually, the GCN part captures stable, smoothed structure, whereas the GAT part introduces attention-driven reweighting of neighbors, and the updated fusion ensures both signals are combined in a controlled way rather than relying on raw concatenation alone.

### 4.5 Hybrid GAT + GraphSAGE

The third hybrid model combines a GAT-based encoder and a GraphSAGE-based encoder applied to the same diagnosis co-occurrence graph, with both components operating directly on the original node features. The GAT component models attention-weighted interactions among neighboring diagnoses, allowing the model to emphasize more informative connections, whereas the GraphSAGE component applies mean aggregation to capture robust local neighborhood statistics inductively. The outputs of the two encoders are normalized to control their relative scales and then fused into a single node representation through a learned combination mechanism, enabling the model to balance attention-driven selectivity with stable neighborhood aggregation. The resulting fused embedding is used for link prediction via the same dot-product (Hadamard-based) decoder as in the other architectures.

### 4.6 Hybrid GCN + GraphSAGE

The second hybrid model applies a GCN-based encoder and a GraphSAGE-based encoder to the same diagnosis co-occurrence graph, with both components operating on the original node features. The GCN component captures globally smoothed structural context through multiple convolution layers, while the GraphSAGE component uses mean aggregation to model localized neighborhood information. The outputs of the two encoders are normalized and then fused into a single node representation using a learned combination mechanism, ensuring that neither global smoothing nor local aggregation dominates the final embedding. The resulting representation is used for link prediction with the same dot-product (Hadamard-based decoder) as above. This design combines complementary inductive biases, allowing the model to exploit both broad graph structure and flexible neighborhood aggregation.

### 4.7 Decoder as the Prediction Module

Given node embeddings **z**_*i*_, **z**_*j*_ ∈ ℝ^*d*^, the decoder computes a logit, followed by a sigmoid function, and predicts co-occurrence likelihood of obstetric and non-obstetric diagnoses.

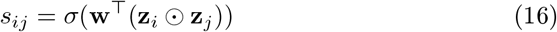

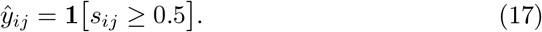

If *ŷ*_*ij*_ = 1, the model considers that obstetric and non-obstetric diagnosis to form a clinically meaningful bridge within pregnancy care. We train the models using binary cross-entropy loss over logits given the ground-truth binary labels *y*_*ij*_ ∈ {0, 1}.

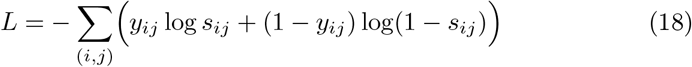

We report area under the receiver operating characteristic curve (AUROC) and area under the precision-recall curve (AUPRC) as primary means of co-occurrence prediction metrics, with F1-score, Matthews correlation coefficient (MCC), accuracy, sensitivity and specificity.

## 5 Results

### 5.1 Overall Performance Analysis

Table 1 summarizes the overall results of all considered models in terms of four well-known performance metric: AUROC, AUPRC, F1, and MCC. We report mean *±* standard deviation over randomly selected 5-fold cross validation sets.

**Table 1.** Obstetric and non-obstetric diagnoses co-occurrences prediction performance across models (mean*±*SD over 5-fold cross-validation).

| Model | AUROC | AUPRC | F1 | MCC |
| --- | --- | --- | --- | --- |
| GCN | 0.835±0.023 | 0.836±0.027 | 0.676±0.006 | 0.134±0.051 |
| GAT | 0.860±0.015 | 0.853±0.019 | 0.790±0.017 | 0.560±0.039 |
| GraphSAGE | 0.889±0.013 | 0.880±0.018 | 0.812±0.010 | 0.621±0.025 |
| GCN+GAT | 0.824±0.011 | 0.810±0.015 | 0.760±0.006 | 0.465±0.017 |
| GAT+GraphSAGE | 0.889±0.013 | 0.883±0.015 | 0.811±0.009 | 0.614±0.020 |
| GCN+GraphSAGE | **0.897±0.012** | **0.894±0.016** | **0.817±0.012** | **0.636±0.025** |

We first attempt to understand the efficacy of the standalone architectures, such as GCN, GAT and GraphSAGE. Among these, GraphSAGE outperforms GCN and GAT in all metrics, particularly by an average of 5% and 2%, respectively, when comparing AUROC. Later hybrid models demonstrate that GraphSAGE is the core component of developing an effective model for predicting obstetric and non-obstetric diagnoses co-occurrences. Additionally, GCN results poor performance among these set of models, especially GCN drops to 0.134 *±* 0.051 of MCC compared with GraphSAGE’s 0.621 *±* 0.025.

Next, we studied the hybrid architectures. GCN+GraphSAGE outperforms all hybrid and standalone methods across all performance metrics. Hybrid GAT +GraphSAGE model performs closely compared with our best performer. We want to highlight that hybrid GCN+GAT performs worst among all considered methods. The strong influence of GraphSAGE on the top-performing hybrid models, as well as the standalone GraphSAGE model, suggests that on this moderately sized and relatively dense diagnosis graph, neighborhood aggregation is particularly effective at capturing the underlying predictive structure. While GAT achieves competitive results, it lags behind GraphSAGE, indicating that attention-based mechanisms alone are somewhat less effective than aggregation-based inductive biases in this setting. In contrast, the standalone GCN model performs the worst among the single-GNN family, suggesting that spectral smoothing alone is not enough to fully utilize the graph structure. Overall, the results support the idea that combining complementary inductive biases-such as GCN-style smoothing with GraphSAGE-style aggregation gives the most reliable prediction of obstetric and non-obstetric diagnoses co-occurrences.

### 5.2 Precision–recall and Receiver-operating Curve analysis

The precision–recall curves and ROC curves in Fig. 1(A,B) provide a more detailed view of ranking quality across thresholds. All models achieve AUPRC values well above the expected baseline for a balanced test set, confirming that they rank true cross-community links above negatives most of the time. Graph-SAGE shows the strongest precision-recall curve among the standalone baselines, maintaining precision above 0.8 across a wide range of recall, followed by GAT, while GCN performs the worst. The hybrid GCN+GraphSAGE achieves the best AUPRC, with a curve that stays close to the upper left corner, meaning it can recover a large fraction of true obstetric and non-obstetric co-occurrence links while retaining high precision. The hybrid GAT+GraphSAGE performs similarly, with only a slight drop in precision at very high recall. In contrast, the standalone GAT curve drops earlier as recall increases, which reflects its difficulty in distinguishing borderline positive edges from negatives.

**Fig. 1.**
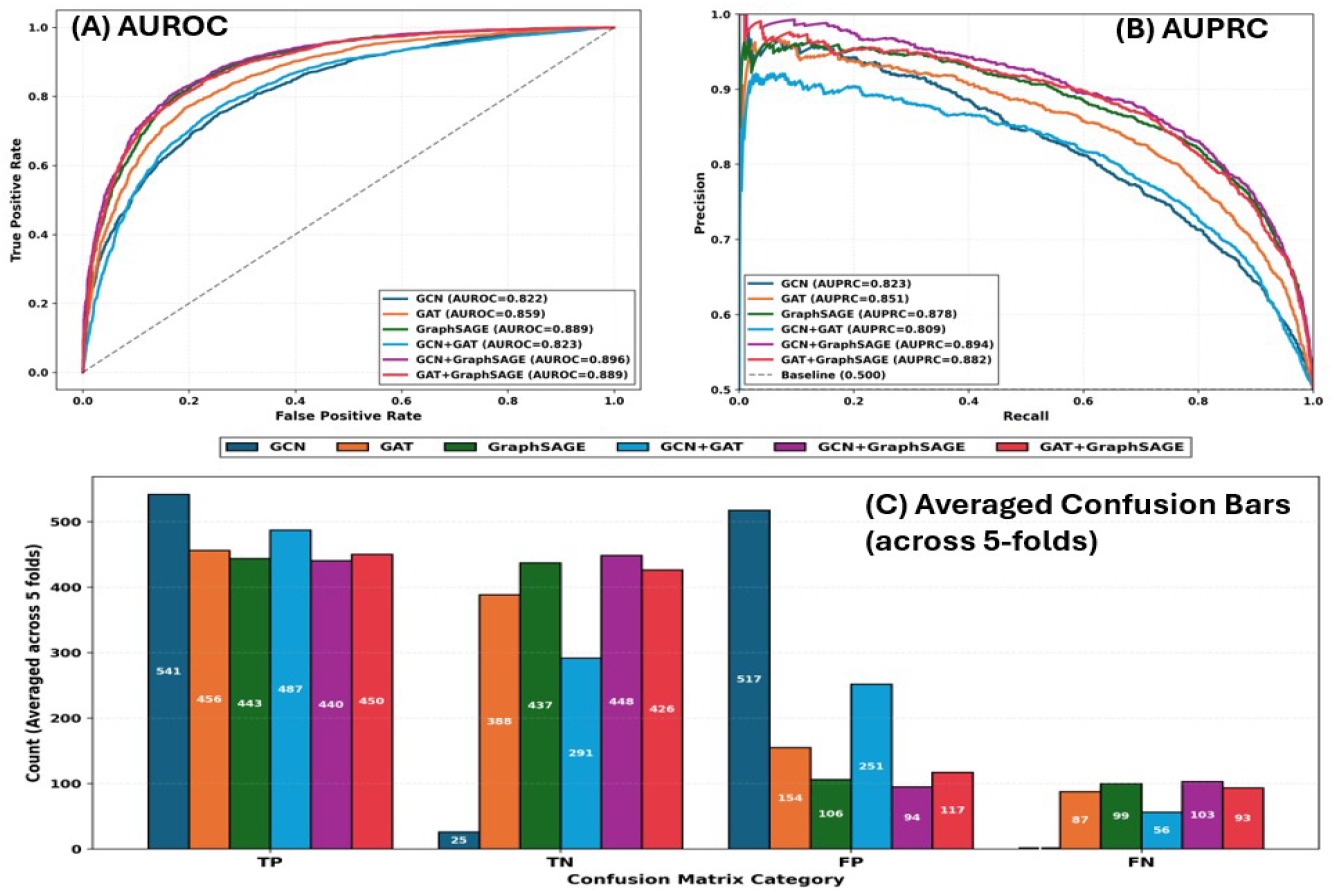
Overall performance analysis of the GNN models. (A) Area under the received operating curve (AUROC) and (B) area under the precision-recall curve (AUPRC) on the test set. (C) Confusion bars averaged over 5-fold cross-validation test sets. TP: true positive, FN: false negative, FP: false positive, FN: false negative.

### 5.3 Error Patterns from Confusion Bars

Figure 1(C) presents averaged confusion bars on the held out test set across 5-fold cross validation. Most models, except GCN, achieved high true negative counts, confirming that they are effective at rejecting non-existent obstetric and non–obstetric diagnoses co-occurrence links. The standalone GCN did not perform well, as it rendered very few true negatives and significantly higher false positives as compared to other models, which aligns with its lower F1 and MCC. The Hybrid GCN+GAT, although not as bad as the standalone GCN model, did render fewer true negatives and more false positives compared to its other better-performing counterparts, which corresponds to its lower F1 and MCC as well. The hybrid GCN+GraphSAGE had the overall best confusion bar pattern as it identified the most true negatives and the fewest false positives, although having slightly higher false negatives as compared to other counterparts. Hybrid GAT+GraphSAGE followed behind, reducing false negatives at the cost of a modest increase in false positives. The pure GAT model displays more false positives but slightly fewer false negatives than the Hybrid GCN+GraphSAGE model. In qualitative terms, the top two hybrid models are better at identifying clinically meaningful bridges without dramatically inflating the number of spurious links, which is important if the predictions are to be used as candidates for risk stratification or hypothesis generation in pregnancy care.

The ROC curves tell a consistent story. All models dominate the diagonal random baseline, but the top two hybrid architectures show the largest area under the curve and stay closest to the upper left corner, indicating a better trade-off between sensitivity and specificity across thresholds. Taken together, the table and plots demonstrate that while GraphSAGE is already a strong single-architecture baseline, GCN performs comparatively worse, and hybrid models that combine aggregation with attention or stacked aggregation deliver the best link prediction performance on the pregnancy diagnosis co-occurrence graph. This curve trend is further supported by the validation AUROC trajectories, where GCN shows an early plateau and limited improvement with continued training, suggesting over-smoothing and reduced discriminative capacity in dense neighborhoods. In contrast, GraphSAGE and GraphSAGE-based hybrids continue to improve and converge to higher AUROC, indicating that explicit neighborhood aggregation better preserves local structural signals, while hybridization mitigates the limitations of spectral smoothing alone.

### 5.4 Top Predicted Obstetric and Non-Obstetric Diagnoses Co-occurrences

To characterize the most salient co-occurrences between obstetric and non-obstetric diagnoses, we applied the best-performing hybrid GCN+GraphSAGE model to all non-existing cross-community pairs, retained the top 10 links per fold according to the predicted edge probabilities, and then aggregated predictions across the 5-folds using a ranking score based on the mean predicted probability multiplied by a recurrence term, mean_probability *×* log(num_folds + 1), which favors edges that both achieve high confidence and consistently appear across multiple folds. Table 2 reports the 10 co-occurrence edges with the highest mean scores. All pairs in the table have mean predicted probabilities above 0.999, which indicates that the model is extremely confident that these diagnosis pairs behave like true co-occurring links in the pregnancy graph, despite not being observed in the original data.

**Table 2.** Top 10 Predicted Cross-Community Diagnosis Associations Between Obstetric and Non-Obstetric Conditions Using hybrid GCN+GraphSAGE Model with 5-fold cross-validation.

| Rank | node 1 name | node 2 name | Mean score |
| --- | --- | --- | --- |
| 1 | Hypothyroidism | Disorder of pregnancy | 0.9999 |
| 2 | High risk pregnancy | Elderly primigravida | 0.9999 |
| 3 | Suspected fetal abnormality affecting management of mother | High risk pregnancy | 0.9999 |
| 4 | Single live birth | Endocrine, nutritional and metabolic disease complicating pregnancy, childbirth and puerperium | 0.9999 |
| 5 | Hypothyroidism | Third trimester pregnancy | 0.9998 |
| 6 | Hypothyroidism | Second trimester pregnancy | 0.9997 |
| 7 | Uterine leiomyoma | Disease of the digestive system complicating pregnancy, childbirth and/or the puerperium | 0.9997 |
| 8 | Third trimester pregnancy | Anemia | 0.9995 |
| 9 | Abnormal findings on antenatal screening of mother | Disorder of pregnancy | 0.9995 |
| 10 | Abnormal findings on antenatal screening of mother | Finding related to pregnancy | 0.9995 |

The top-ranked connections highlight clinically plausible bridges involving obstetric complications and systemic medical conditions. Many of the highest scoring pairs link endocrine and metabolic disorders, particularly “hypothyroidism” and “type 2 diabetes mellitus without complication”, to trimester-specific pregnancy diagnoses and broader obstetric risk labels such as “disorder of pregnancy” and “high-risk pregnancy”. These links reflect the fact that chronic conditions are frequently documented along pregnancy-related stage or risk stratification codes, rather than being encoded as a single combined diagnosis.

Several additional high-scoring connections associate pregnancy stage codes with “anemia”, “maternal and/or fetal conditions affecting labor and delivery”, and other pregnancy-related findings. From a clinical perspective, these predictions suggest that the model is capturing recurring documentation patterns in which systemic conditions and obstetric status are recorded in parallel across encounters, even when their interaction is implied rather than explicitly specified. In the context of our research question, the table provides concrete candidate bridges where non-obstetric diagnoses may coincide with pregnancy progression or delivery-related risk, and it highlights specific diagnosis pairs that could warrant targeted chart review and longitudinal analysis in future obstetric risk stratification studies.

## 6 Conclusion

In this work, we formulate the obstetric and non-obstetric diagnosis co-occurrence in pregnancy as a link prediction problem on a diagnosis graph and show that GNNs can accurately rank clinically meaningful comorbidity links from noise. Using a homogeneous diagnosis co-occurrence graph enriched with structural and demographic node features, most models achieved strong discrimination between true and negative cross-community pairs except the standalone GCN model. Particularly, hybrid GCN+GraphSAGE and GAT+GraphSAGE encoders reach the highest performance and identify top-ranked links that align with endocrine, metabolic, and hematologic conditions co-occurring with pregnancy stage and risk-related diagnoses, catching clinically plausible patterns that are often under-recorded within single visits. These results suggest that graph-based link prediction is a useful tool to generate prioritized lists of obstetric and non-obstetric diagnosis pairs that may influence pregnancy trajectories and postpartum recovery.

In downstream clinical workflows, the proposed model could serve as a retrospective screening and prioritization layer rather than a diagnostic tool. Predicted high-confidence pairs of obstetric-non-obstetric diagnosis may be used to identify comorbidity patterns for targeted chart review, guide risk-stratified monitoring during prenatal care, or inform the design of rule-based alerts within existing EHR systems. Importantly, such integration would occur offline or asynchronously, supporting clinicians by detecting plausible diagnosis associations rather than generating automated clinical decisions.

This study has several limitations that open up directions for future work. Our experiments rely on a single sampled EHR dataset, focus exclusively on diagnosis codes within pregnancy-related encounters, and treat co-occurrence within a visit as the only notion of relatedness, without modeling temporal order, severity, treatment, or patient-level outcomes. The link predictions are observational and should be interpreted as hypotheses rather than evidence of causal relationships, fairness across patient subgroups, or prospective utility at the point of care. Future work should validate these findings in larger and more diverse cohorts, incorporate medications, procedures, and laboratory results into heterogeneous or temporal graphs, explore graph transformers and uncertainty-aware models, and connect diagnosis-level link prediction to patient-level risk scores that could be evaluated in collaboration with clinicians in real obstetric surveillance workflows.

## Footnotes

3 This work has been accepted at the 18th International Conference on Bioinformatics and Computational Biology (BICOB’26)

